# On the estimation errors of *K_M_* and *V* from time-course experiments using the Michaelis–Menten equation

**DOI:** 10.1101/068015

**Authors:** Wylie Stroberg, Santiago Schnell

## Abstract

The conditions under which the Michaelis–Menten equation accurately captures the steady-state kinetics of a simple enzyme-catalyzed reaction is contrasted with the conditions under which the same equation can be used to estimate parameters, *K_M_* and *V*, from progress curve data. Validity of the underlying assumptions leading to the Michaelis–Menten equation are shown to be necessary, but not sufficient to guarantee accurate estimation of *K_M_* and *V*. Detailed error analysis and numerical “experiments” show the required experimental conditions for the independent estimation of both *K_M_* and *V* from progress curves. A timescale, *t_Q_*, measuring the portion of the time course over which the progress curve exhibits substantial curvature provides a novel criterion for accurate estimation of *K_M_* and *V* from a progress curve experiment. It is found that, if the initial substrate concentration is of the same order of magnitude as *K_M_*, the estimated values of the *K_M_* and *V* will correspond to their true values calculated from the microscopic rate constants of the corresponding mass-action system, only so long as the initial enzyme concentration is less than *K_M_*.

## 1. Introduction

The fundamental equation of enzyme kinetics is the Michaelis–Menten (MM) equation, which relates the rate of an enzyme-catalyzed reaction to the concentration of substrate [1, 2]. The MM equation is typically derived using the steady-state assumption as proposed by Briggs and Haldane [3]. It is characterized by two parameters: the Michaelis constant, *K_M_*, which acts as an apparent dissociation constant under the assumption of steady-state, and the limiting rate, *V* (or the catalytic constant, *k_cat_* if the enzyme concentration is known) [4]. These parameters are often viewed as thermodynamic properties of an enzyme–substrate pair, and hence depend on conditions such as pH and temperature, but not on time-dependent enzyme nor substrate concentrations [5]. As a result, measuring *K_M_* and *V* are essential to characterizing enzymatic reactions [6]. However, the treatment of *K_M_* and *V* as constants with respect to enzyme and substrate concentrations relies on simplifying assumptions relating to the quasi-steady-state of the intermediate complex formed by the enzyme and substrate [7]. If conditions for the reaction lie outside the range for which the simplifying assumptions are valid, *K_M_* becomes dependent on the concentration of the substrate, and hence, on time. Experiments to estimate *K_M_* must be conducted under conditions for which the MM equation is valid [7, 8]. This can be problematic since it is generally necessary to know *K_M_* a priori in order to insure the experimental conditions meet the requirements for the using MM equation. Additionally, values of *K_M_* and *V* measured under valid experimental conditions can only be transferred to cases that also meet the requirements. Since this is often not the case in vivo, using values of *K_M_* and *V* measured in vitro to predict the activity of an enzyme in living organisms can often be seriously unreliable [9].

The range of substrate and enzyme concentrations over which the MM equation can be applied has long history of theoretical investigation [see 8, for a recent review], and requires two assumptions be valid. The first, called the steady-state assumption, implies that the timescale for the formation of the intermediate complex is much faster than that of the conversion of the substrate into product [10]. The second, called the reactant-stationary assumption, implies that the fast, transient period in which the steady-state population of intermediate complex first forms, depletes only a negligible amount of substrate [11]. It has been shown that the reactant-stationary assumption is more restrictive and, if valid, the reaction velocity (after the initial transient period) will follow the MM equation and be well-characterized by the parameters *K_M_* and *V* [10, 12, 8].

At first sight, it is tempting to assume that, when the MM equation is valid, experimental data should also yield accurate estimates of *K_M_* and *V* [13, 14]. However, the conditions for the validity of the steady-state and reactant-stationary assumptions are based on a forward problem, i.e. one in which the parameters are known. Estimating parameters from experimental data, on the other hand, is an inverse problem [15]. Extracting true values of parameters from data requires a stable and unique inverse mapping that is not guaranteed by the existence of a solution to the forward problem [see 16, for example]). Hence, even in cases where the assumptions underlying the MM equation are valid, and the MM equation accurately fits an experimental progress curve, the values of *K_M_* and *V* estimated from the data may differ significantly from their true values.

Understanding the conditions for which the inverse problem is well posed is crucial for the effective and efficient design of experiments. When designing enzyme progress-curve experiments, one typically must choose the initial concentrations of the substrate and enzyme (although the enzyme concentration may not always be adjustable), as well as the time span and sampling frequency for data collection [17]. Hence, useful experiments require conditions that both satisfy the conditions for which MM kinetics are to be expected, and lead to the most informative set of data for constraining parameter values. Early use of progress curves to determine kinetic parameters focused on linearization of the rate equations or efficient integration and optimization algorithms for fitting parameters [18, 19, 20, 21, 22]. As these algorithms evolved, computational tools for analysis of progress curve data increased the accessibility and popularity of progress curve experiments [23, 24, 25]. However, less attention has been paid to the design of progress curve experiments. Initial research applied sensitivity analysis [26], and information-theoretic approaches [27] to estimate optimal initial substrate concentrations and the most sensitive portion of the progress curve, and hence, the most useful portion for parameter estimation. Vandenberg et al. [26] found that the largest feasible substrate concentration and the section of the progress curve for which the substrate concentration is between 60–80% of the initial value maximized the sensitivity of the fitted parameters. However, maximizing the sensitivity of the data collection range does not necessarily guarantee minimization of the errors in the fitted parameters. To address this, Duggleby and Clarke [17] assessed the optimal initial substrate concentration and data spacing under the criterion of minimal standard error of *K_M_*. The optimal design of Duggleby and Clarke differs from that of Vandenberg et al. in that an initial substrate concentration 2 to 3 times *K_M_* is recommended. It was also found that data should be collected until the extent of the reaction is 90%. These recommendations have become the de facto “rule of thumb” for progress curve experimental design. In determining these recommendations, the authors evaluated their parameter estimates in comparison to parameter values obtained through initial rate experiments on the same enzymatic systems, and to simulated progress curves calculated by integrating the MM equation and adding random fluctuations. Hence, no connection was made to the underlying microscopic rate constants describing the mass-action kinetics of the systems, meaning the accuracy of the estimates could not be assessed relative to the “true” values of *K_M_* and *V* as defined in terms of microscopic rate constants. A similar approach was later taken to evaluate the capacity of a closed-form solution to the MM equation to fit progress curves [28].

The work of Duggleby and colleagues provide guidance for when the parameters in the MM equation, *K_M_* and *V*, are most robustly estimated from progress curve experiments, but do not assess whether the fitted parameters are the same as those defined in terms of microscopic rate constants. With improved fitting algorithms and greater computational power, interest has grown in the direct determination of microscopic rate constants through fitting of progress curves with numerically-integrated rate equations [29]. Although appealing, this approach can only provide accurate estimates for parameters that are sensitive to the given experimental conditions. Under experimental conditions for which the mass-action rate equations reduce to the MM equation, this procedure will necessarily lead to overfitting. Designing experiments from which *K_M_* and *V* can be unambiguously determined requires assessing the experimental conditions in terms of the requirements for the validity and uniqueness of the MM equation. Moreover, given the massive amounts of data generated by the biomedical science community, scientists must be cognizant of the strengths and weakness of quantitative approaches in order to guarantee the reproducibility of published research data.

In this work, we seek to address the issue of estimating parameters from progress curves of single-substrate, single-enzyme-catalyzed reactions quantitatively. In Section 2, we review the validity of the steady-state and reactant-stationary assumptions, and quantify errors incurred by making these assumptions. In Section 3, we discuss the inverse problem associated with estimating parameters based on the MM equation. In doing so, we derive a new condition based on time-scale separation of the linear and nonlinear portions of the progress curve that indicates when both *K_M_* and *V* can be estimated from a single experiment. Numerical experiments are then conducted in Section 4 to verify and quantify the range of experimental conditions that allow for veracious estimations of*K_M_* and *V*. We conclude with a discussion of the results in Section 5.

## 2. The forward problem: the Michaelis–Menten equation and the conditions for its validity

In the simplest, single-enzyme and single-substrate reaction, the enzyme *E* reacts with the substrate *S* to form and intermediate complex *C*, which then, under the action of the enzyme, forms a product *P* and releases the enzyme,

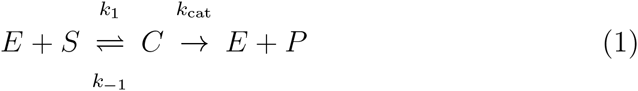

where *k*_1_ and *k*_−1_ are microscopic rate constants, and *k*_cat_ is the catalytic constant [4]. Applying the law of mass action to reaction mechanism (1) yields four rate equations

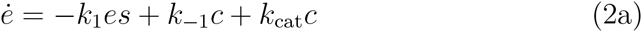

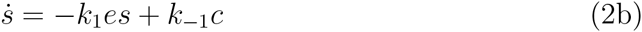

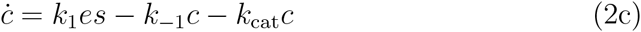

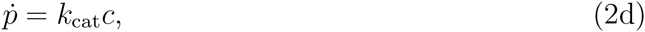

where lowercase letters represent concentrations of the corresponding uppercase species. Typically, in test tube enzyme binding assays the initial conditions are taken to be

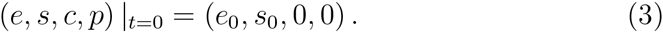

Additionally, the system obeys two conservation laws, the enzyme and substrate conservation laws,

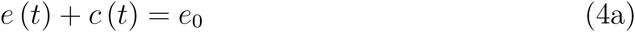

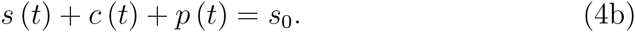

Using (4a) to decouple the enzyme concentrations, the redundancies in the system (2) are eliminated to yield

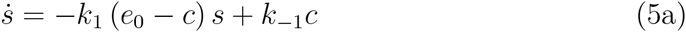

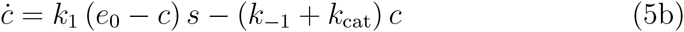

where *e*(*t*) and *p*(*t*) are readily calculated once *s*(*t*) and *c*(*t*) are known. If, after an initial, rapid buildup of *c*, the rate of depletion of *c* approximately equals its rate of formation, *c* is assumed to be in a quasi-steady state [3], i.e.

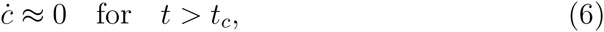

where *t_c_* is the timescale associated with the initial transient buildup of *c* [10]. The steady-state assumption (6), in combination with (5), leads to

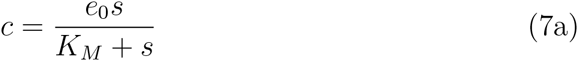

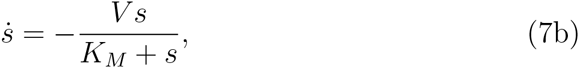

where *V* = *k*_cat_*e*_0_ and *K_M_* = (*k*_−1_ + *k*_cat_)/*k*_1_. Hence, the system (2) is reduced to an algebraic-differential equation systems with one single differential equation for *s*. However, since (7) is only valid after the initial transient time period, *t_c_*, a boundary condition for *s* at *t* = *t_c_* must be supplied. To do this, it is assumed that very little substrate is consumed during the initial transient period (the reactant-stationary assumption) such that

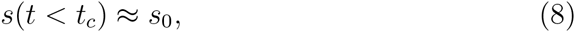

which provides an initial condition for (5a) under the variable transformation *t* → *t*–*t_c_*. Substituting (7a) into (2d), one obtains, the rate of the reaction (1)

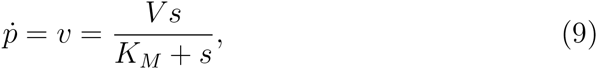

relating the rate of product formation to the substrate concentration. Equation (9) is the MM equation, and the system of equations (7a), (7b), and (9) govern the dynamics the complex, substrate, and product, respectively, under the steady-state assumption.

The conditions under which the steady-state assumption (6) and reactant-stationary assumption (8) are valid have been extensively studied. Segel [10] showed that the steady-state assumption is valid so long as

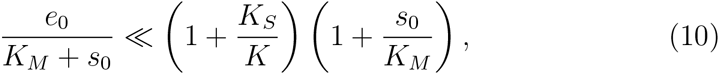

where *K_S_* = *k*_−1_/*k*_1_, and *K* = *k*_cat_/*k*_1_. For the reactant-stationary assumption to be valid, they derived the condition

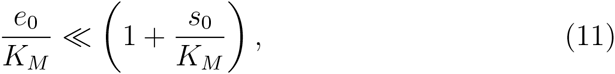

which is more stringent than condition (10), and hence dictates the conditions under which the MM equation can be applied. Interestingly, it has been shown that condition (11) is independent from (10) for several enzyme catalyzed reactions [11].

### 2.1. Quantitative analysis of the errors induced by the steady-state and reactant-stationary assumptions

To gain a quantitative understanding of the inequalities expressed in (10) and (11), an accurate assessment of the difference between the solution to system (5) and the reduced equations (7) is required. For our analysis, we compare progress curves of the substrate calculated with numerical solutions to the exact law of mass action system (5a) and the reduced equation (7b) under the steady-state assumption. Note that the reduced rate equation (7b) is effectively the MM equation for the substrate depletion. The *concentration error* as a function of time is calculated as

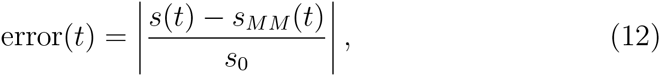

where *s*_MM_ is the substrate concentration calculated using the reduced equation (7b) and | · | denotes the absolute value. To form a scalar measure of the error, we use the maximal value of the concentration error over the time course of the reaction. Contours of the maximum concentration error in the plane of initial enzyme and substrate concentrations (normalized by *K_M_*) are shown in **Fig. 1**. Additionally, conditions (10) and (11) are plotted for the cases when the right-hand sides are ten times the left-hand sides to represent the much less condition numerically. For all values of *κ* = *k*_−1_/*k*_cat_ = *K_S_*/*K*, condition (11) is sufficient to guarantee small errors when using the MM equation. However, **Fig. 1A** shows that when *κ* is small – implying the reverse step in reaction (1) is negligible – small values of *s*_0_/*K_M_* yield small errors, regardless of the initial enzyme concentration.

**Figure 1:**
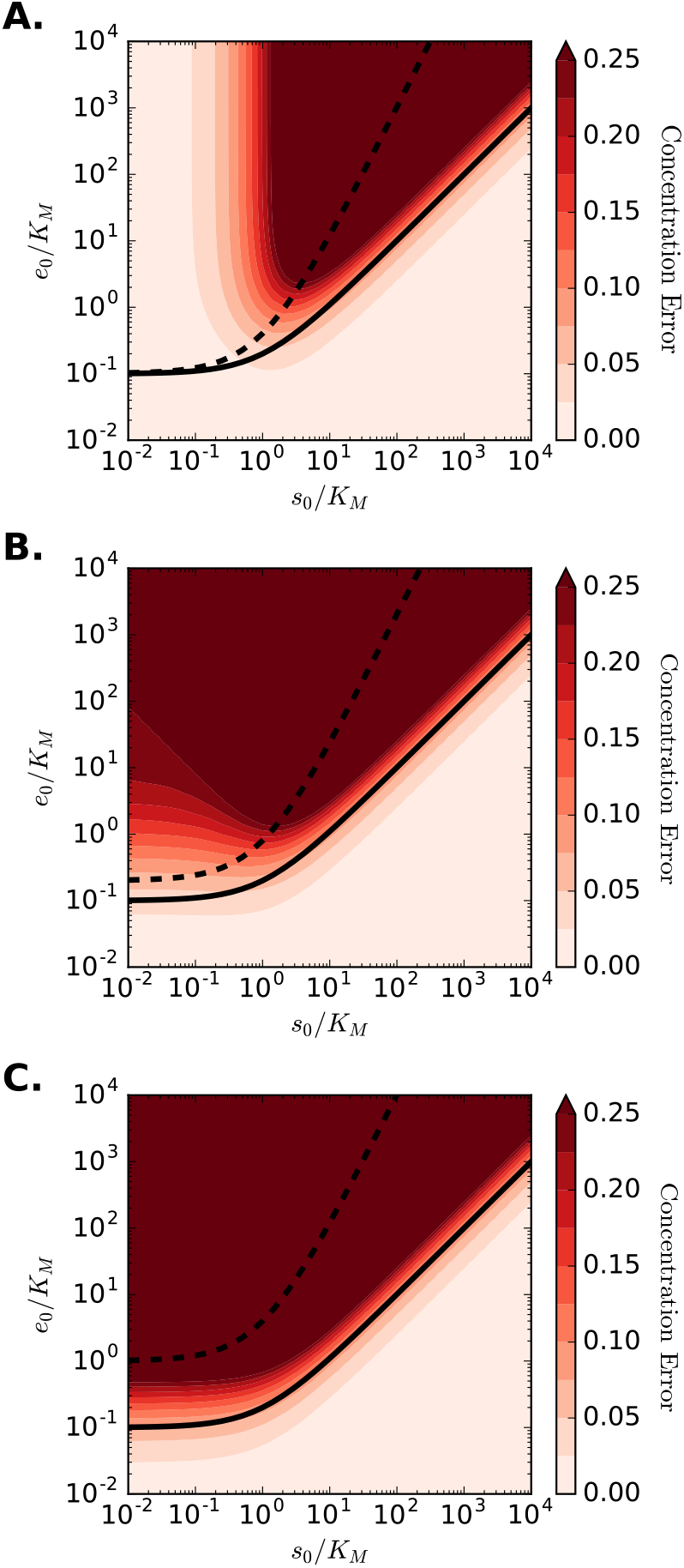
Concentration error contours in the *e*_0_/*K_M_*–*s*_0_/*K_M_* plane. The maximal concentration errors are plotted in the plane of initial enzyme and substrate concentrations, normalized by *K_M_*. The dashed black line corresponds to the condition for steady-state assumption (10), while the solid black line corresponds to the reactant stationary condition (11). Each panel shows different values of *K_S_* and *K*, while *K_M_* = 1 for all cases. Panel **A**: *K_s_* = 0.1, *K* = 0.9; Panel **B**: *K_S_* = 0.5, *K* = 0.5; Panel **C**: *K_S_* = 0.9, *K* = 0.1.

The observed errors can be understood by considering the influence of small *κ* and *s*_0_/*K_M_* on the system (5). When *k* ≪ 1, reaction (1) strongly favors the production for *P* from *C* as opposed to the disassociation of *C* back to *E* and *S*. This reduces the reaction mechanism (1) to the van Slyke–Cullen mechanism [30] as *K_M_* & *K*. The requirement *s*_0_/*K_M_* ≫ 1 implies that the formation of *C* is slow compared to the formation *P* and the disassociation of *C*. Taken together, these two requirements provide an ordering of timescales such that the formation of *C* is slow compared to the action of the enzyme to form *P*, but fast compared to the disassociation of the intermediate complex, effectively reducing the rate equation for the substrate depletion (5a) to *ṡ* ≈ −*k*_1_*e*_0_*s*. Similarly, under the same condition, the MM equation for substrate (7b) reduces to the same expression. Hence, under these conditions, the MM equation accurately represent the system dynamics, even though condition (11) is violated.

The condition for the validity of the reactant-stationary assumption (11) is a sufficient condition for the MM equation to be valid. In essence, this says that for a known set of parameter values, if the reactant-stationary assumption is valid, the dynamics of the reduced system (7) will closely approximate the dynamics of the full system (5). However, the MM equation is often used to estimate *K_M_* and *V* from experimental data, which requires solving an inverse problem. Solutions to the forward problem do not guarantee the existence or uniqueness of the inverse problem, hence it is not clear that the conditions for the validity of the reduced forward problem correspond to the conditions required to accurately estimate rate constants. This issue is investigated in the following section.

## 3. The inverse problem: Estimation of *K_M_* and *V*

The experimental estimation of the parameters *K_M_* and *V* is used to characterize enzyme-catalyzed reactions. In general, *K_M_* and *V* can be estimated through either initial rate experiments [see 31, for a recent review] or direct analysis of time course data [28]. In initial rate experiments, a series of enzyme assays with differing substrate concentrations are performed and initial reaction rates are calculated from the linear portion of the progress curve (after the initial fast transient, *t_c_*, and before substrate depletion becomes influential). The MM equation for either substrate or product is then fit to the initial rates as a function of initial substrate concentration, yielding *K_M_* and *V*. When time course data is used, the integrated implicit [32] or closed-form [33] of the MM equations are fit directly to time series through nonlinear regression, providing estimates for *K_M_* and *V*. Although initial rate experiments are more commonly used, they require numerous assays with different substrate concentrations to determine *K_M_* and *V*. Alternatively, time course analyses have the advantage that *K_M_* and *V* can be estimated from a single experiment, making them potentially much cheaper when expensive I reactants are required, and less time consuming [34, 35, 36, 7]. Hence, in I this work, we consider the problem of parameter estimation directly from progress curves, specifically, those for the concentration of substrate.

Inverse problems are typically formulated in terms of an operator, *F*, mapping the space of parameters, *Q*, to the space of observations, *Y*, i.e.

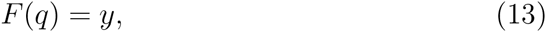

where *q* ∈ *Q* is a vector of parameters, and *y* ∈ *Y* is a vector of observed quantities. In general, *F* = *G* o *H* is a composite of the solution operator *S*, which maps a parameter vector *q* to a solution vector 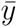 of the underlying ordinary differential equation for the rate equations, and and the observation operator *R*, which takes 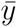 to the observable *y* [37]. For example, if fluorescent markers are used to tag substrate molecules, and fluorescent intensity is measured at times *t_i_, G* is then the mapping between the fluorescent intensity at times *t_i_* and substrate concentration, and *H* is the solution to the rate equations (7). *G* effectively samples the solution to the rate equation model at the observation times and converts those concentrations to the experimental observables.

For the present study, we assume the concentrations are observed directly, hence *G* is simply a sampling of the integrated rate equations (5). Specifically, we consider the case in which the concentration of the substrate is measured at discrete times *t_i_* and *H* is the solution to (7). The inverse problem consists of finding a parameter vector *q* solving (13). However, (13) is generally ill-posed due to experimental noise. Even in the absence of experimental error, the inverse problem will be ill-posed, because the MM equation only approximates the mass action rate equations (5), even when the steady-state and reactant-stationary assumptions are valid. The exact inverse problem must then be reformulated as a least-squares optimization problem to minimize the function

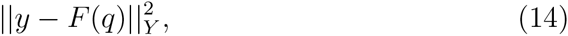

where ║ · ║_*Y*_ is the *L*_2_ norm on *Y*. The sensitivity of (14) to changes in parameter values is measured by the local condition number for the first order optimality condition. The condition number is given by the ratio of the maximum and minimum eigenvalues of the matrix

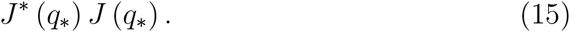

In the above expression, *J* is the Jacobian of the mapping *F*, *q*_*_ is the “true” parameter vector and *J*^*^ denotes the conjugate transpose of *J*. Ill-conditioning implies small errors in the data (or model) can result in large errors in the estimated parameters. Although many features of a problem can affect the conditioning (such as proper choice of units) [38], of particular importance when fitting the MM equation is the correlation of the parameters. When the parameters are highly correlated the model is incapable of uniquely determining the parameters because, as the correlation coefficient tends toward 1, the parameters become linearly dependent. In this case, at least one column of *J* will be approximately a linear combination of the others, and hence not invertible.

Effectively, this dictates when the mass action model (5), which depends on three parameters (*k*_1_, *k*_−1_, *k*_cat_), reduces to the MM model (7) with parameters (*K_M_, V*). Under experimental conditions for which the reactant stationary assumption is valid, it is not possible to estimate all three rate constants from the mass action model using time course data. Similarly, within the region of validity for the reactant stationary assumption, there are sub-regions in which columns of *J* become nearly linearly dependent, and hence prohibit estimation of both *K_M_* and *V* from time course data. To see where this rank deficiency occurs we consider two regions in the *s*_0_/*K_M_*–*e*_0_/*K_M_* plane. In both, the conditions for the validity of the reactant stationary assumption are met. Additionally, in the first case *s*_0_ ≪ *K_M_*. Since *s* < *s*_0_ for all *t*, we can expand (5a) in powers of *s*/*K_M_*. Truncating this expansion at order two leads to

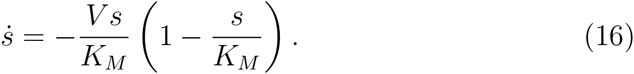

To lowest order, *ṡ* depends only on the ratio of *V* to *K_M_*, and hence the inverse problem of finding both parameters from time course data will become extremely ill-conditioned at small substrate concentrations (see, **Fig. 2A**).

**Figure 2:**
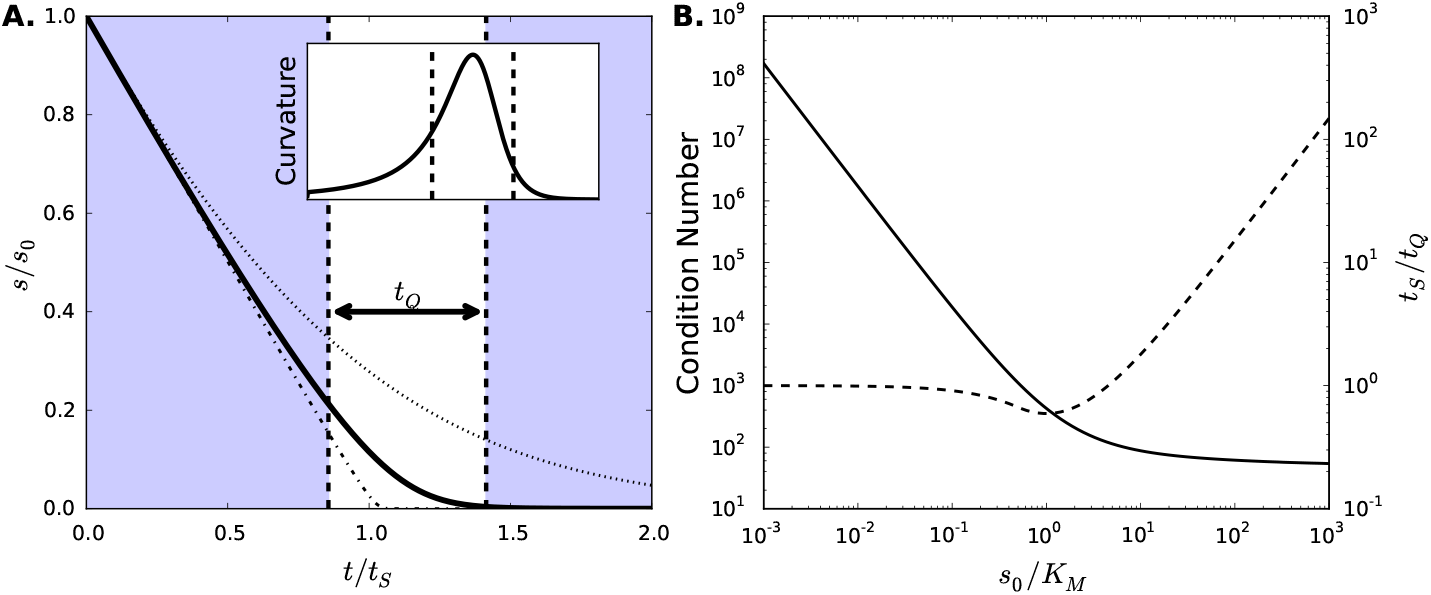
**(A) Substrate progress curves for high, intermediate and low initial substrate concentrations.** Substrate concentrations for differing values of *s*_0_/*K_M_* are plotted as a function of time. When the initial substrate concentration is large (*s*_0_ = 100*K_M_*, dot-dashed line) the substrate depletion is linear until nearly all substrate has been depleted. With low initial substrate concentration (*s*_0_ = *K_M_*, dotted line), the depletion follows a simple exponential. At intermediate values (*s*_0_ = 10*K_M_*, bold solid line), the concentration follows the full hyperbolic rate law and both *K_M_* and *V* can be uniquely identified through regression. The non-shaded region marks the timescale *t_Q_* for the bold curve, centered at the point of maximal curvature for the time course. Inset shows the curvature of the bold progress curve as a function of time, with the *t_Q_*-region demarcated by dashed lines. Parameters for the case shown are: (*k*_1_, *k*_−1_, *k*_cat_) = (1.0,0.5, 0.5), *s*_0_ = 10*K_M_*, *e*_0_ = *K_M_*. **(B) Condition number (solid line) and timescale ratio** *t_S_*/*t_Q_* **(dashed line) as functions of the** *s*_0_/*K_M_*. At small values of *s*_0_/*K_M_*, the inverse problem becomes ill-conditioned. At large values of *s*_0_/*K_M_*, the region of the progress curve providing information about *K_M_* becomes increasingly small.

Next, consider the case in which the substrate is in great excess, i.e. *s*_0_ ≫ *K_M_*. Initially, *s* ≈ *s*_0_, allowing for an expansion of *S* in powers of *K_M_*/*s*_0_, which, to second order, gives

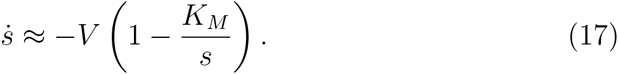

Hence, so long as *s* ≫ *K_M_*, the substrate concentration will decrease linearly with rate –*V*. Eventually, the progress curve must deviate from the initial linearity, and presumably, this curvature should contain information about *K_M_*, allowing for both parameters to be estimated. However, if the time over which the progress curve is nonlinear is small, or equivalently, the initial linear regime very nearly approaches substrate depletion, parameter estimation will fail. Large *s*_0_/*K_M_* can be shown to imply this by comparing the timescale for significant substrate depletion, *t_S_*, with the timescale of high curvature, *t_Q_*. The substrate depletion timescale is given by [10]

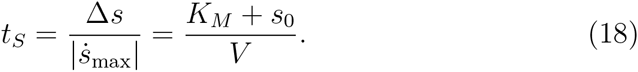

The high-curvature timescale can be estimated with the aid of the second derivative of the substrate concentration,

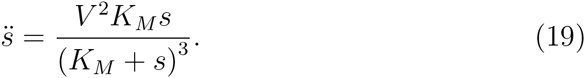

*t_Q_* is defined as the ratio of the total change in velocity of the reaction to the maximum acceleration. The maximum acceleration, found by equating (19) with zero, occurs when *s* = *K_M_*/2 for *s*_0_ ≥ *K_M_*/2, and *s*_0_ otherwise. Since the present analysis concerns high *s*_0_/*K_M_*, the high-curvature timescale is given by

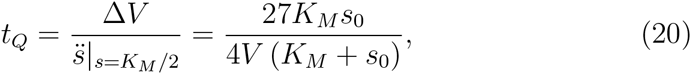

where Δ*V* is the change in reaction velocity through the region of curvature and is equal to *V*. As shown in **Fig. 2A**, *t_Q_* measures the time over which the progress curve has significant curvature. Estimation of parameters from time course data will not be possible when *t_Q_* ≪ *t_S_*, or, upon substitution of (18) and (20), when

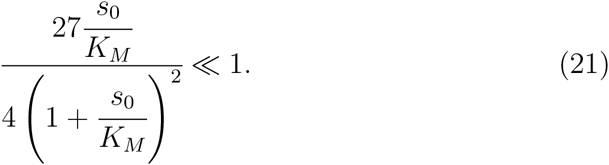

Therefore, as the initial substrate concentration is increased, the proportion of the time course that can yield information about *K_M_* decreases, and measurements will require greater resolution in both time and concentration. **Fig. 2B** shows the condition number and the ratio of the substrate depletion timescale to the high-curvature timescale for a large range of *s*_0_/*K_M_*. At small values of *s*_0_/*K_M_*, ill-conditioning makes parameter extraction intractable, while at large *s*_0_/*K_M_*, measurements must be increasingly precise. Thus, substrate concentrations close to *K_M_* are desirable when determining parameters.

## 4. Numerical experiments

To demonstrate and quantify the regions in which the conditioning of the inverse problem is poor, and the necessary measurements become intractable, we present a systematic numerical analysis of progress curve experiments in this section.

### 4.1. Methodology for numerical progress curve experiments

Numerical experiments consist of first generating progress curve data from the mass action rate equations with a known set of rate constants. Then, the values of *K_M_* and *V* corresponding to those rate constants are estimated by fitting the MM equations to the progress curve. To generate experimental progress curves it is necessary to choose a set of rate constants (*k*_1_, *k*_−1_, *k*_cat_), and experimental protocol. The experimental protocol consists of defining initial conditions, (*s*_0_, *e*_0_, *c*_0_, *p*_0_), a time span for the experimental observation, *t*_obs_, and a sampling frequency *ω*. The system of equations (5) are integrated numerically from *t* ∈ [0, *t*_obs_] and substrate concentrations are recorded every *ω*^−1^ time units, leading to *t*_obs_*ω* data points {*s_i_* (*t_i_*)}.

The data is then fitted using the numerically integrated form of (5a). The nonlinear regression used to calculate the parameters (*K_M_, V*) is performed using the Levenberg-Marquardt algorithm as implemented in SciPy (version 0.17.1, http://www.scipy.org). In many cases, supplying good initial conditions for the optimizer used for the regression is crucial to finding accurate parameter estimates. Since, in actual experiments the values of *K_M_* and *V* are not known a priori, we attempt to roughly estimate their values from the time course data to provide initial conditions for the optimization. To do this, {*s_i_* (*t_i_*)} is differentiated numerically by central differences to give approximate rates {*ṡ_i_*(*t_i_*)}. Then, using (5a), data at any two time points, *t_i_* and *t_j_* can be used to estimate the parameters through

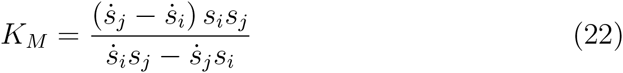

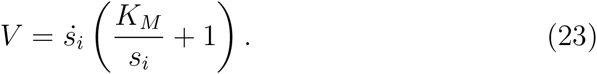

In theory, any two points can be used to estimate *K_M_* and *V*, however, it is best to use data for which the velocity is changing at that greatest rate. Hence, we additionally numerically calculate 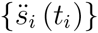 and choose the times directly on either side of the maximum to substitute into (22) and (23). To avoid using data points in the transient regime before the system reaches a quasi-steady state, we consider only the regime for which *s*(*t_i_*) < *s*_0_/2. For actual experiments, noise can make calculations of derivatives subject to large errors, hence smoothing techniques must be used. Additionally, numerous pairs of data points can be used to generate a distribution of estimates, which can then be averaged to give initial conditions for the optimization, similar to [39]. Once the initial conditions for the optimization routine are established, the best-fit values of *K_M_* and *V* can be systematically estimated.

We note that when experimental conditions do not lie in a region for which the reactant stationary assumption is valid, the above technique will naturally provide poor estimates for *K_M_* and *V*. In these regions, we have also used the true values *K_M_* and *V*, calculated from the known rate constants, as initial conditions. Both methods provide qualitatively similar results throughout the regions of parameter space investigated here, and quantitatively agree in the region for which the reactant stationary assumption is valid.

### 4.2. Errors in parameter estimates can be large even when the reactant stationary assumption is valid

Despite the validity of the reactant stationary assumption being sufficient for the MM equation to closely align with the solution to the mass action governing equations, the inverse problem does not provide accurate estimates for parameters within the same range. **Fig. 3** shows errors in estimates of *K_M_* and *V* for a wide range of *e*_0_/*K_M_* and *s*_0_/*K_M_*. Below and above the range plotted for *s*_0_/*K_M_*, the solutions become numerically unstable due to the conditioning problems discussed in Section 3. It is clear however, that even within the range defined by large and small values of *s*_0_/*K_M_*, significant errors are present. At high *s*_0_/*K_M_* and *e*_0_/*K_M_, V* can be accurately determined, but *K_M_* begins to show significant deviation. This is anticipated from the high-*s*_0_/*K_M_* approximation of the substrate rate equation, which depends only on *V*. Additionally, when *s*_0_/*K_M_* < 1, the error contours follow a line for which *e*_0_ ≈ *s*_0_. The condition that enzyme concentration is small relative to that of the substrate was one of the earliest conditions for the validity of the MM equations derived from singular perturbation theory [40]. For the forward problem, Segel [10] showed this condition to be overly restrictive, yet it appears to be appropriate for the inverse problem.

**Figure 3:**
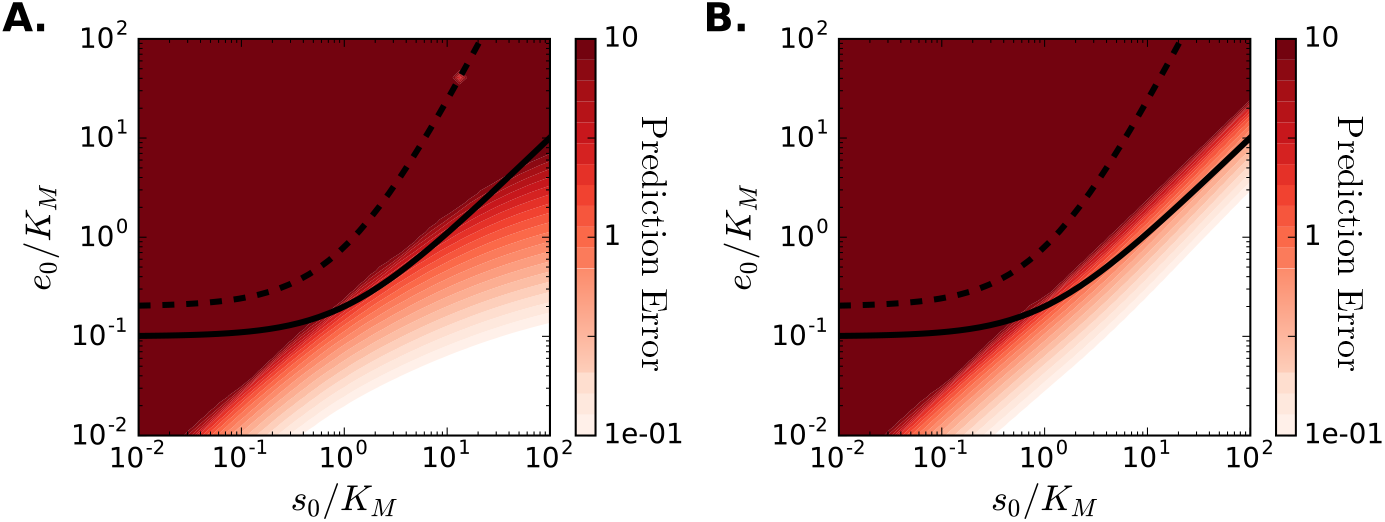
Error contours of the estimated values *K_M_* and *V*. Errors in the predicted values of *K_M_* and *V* for different initial substrate and enzyme concentrations are shown to deviate from the conditions for the validity of the reactant stationary assumption (shown as the solid line). The dashed line corresponds to condition (10). For *s*_0_/*K_M_* values lower than 10^−2^ and greater than 10^2^, the fitting algorithm becomes unstable. Note that the color bar scale is logarithmic, showing errors can be significant. In this figure, *K_S_* = *K* = 10.

An explanation for the condition *e*_0_ ≪ s_0_ can be found by comparing the integrated form of the MM substrate equation with an exponential progress curve that is the limiting solution to the MM equations as *s*_0_/*K_M_* approaches zero. The integrated closed-form of (7b), known as the Schnell-Mendoza equation [33], can be written explicitly in terms of the Lambert-W function [41]

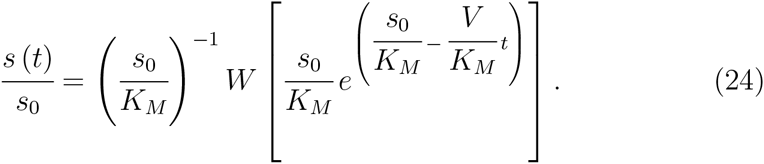

Expanding the above expression about zero and truncating at first order leads to

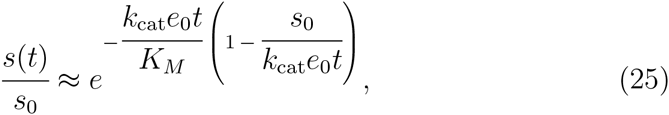

where we have used the definition of *V* to explicitly show the dependence on *e*_0_. The exponential solution takes the form

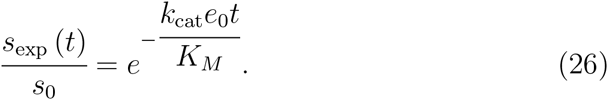

Comparing (25) and (26) shows that the correction provided by the MM solution over the exponential progress curve becomes decreasingly significant as *e*_0_/*s*_0_ becomes large. **Fig 4A** compares the mean concentration errors between the best-fit solutions and the “true” solutions for both the MM equation and an exponential fit. At small values of *e*_0_/*s*_0_, the MM equation provides a distinctly better fit than the exponential solution, allowing both *K_M_* and *V* to be estimated from a single progress curve. As *e*_0_/*s*_0_ increases, the two fitting functions eventually provide the identical fits. This corresponds to an exponential increase in the variance of the estimated parameters (**Fig. 4B**), and indicates that only the ratio *V*/*K_M_* can be determined in this range.

**Figure 4:**
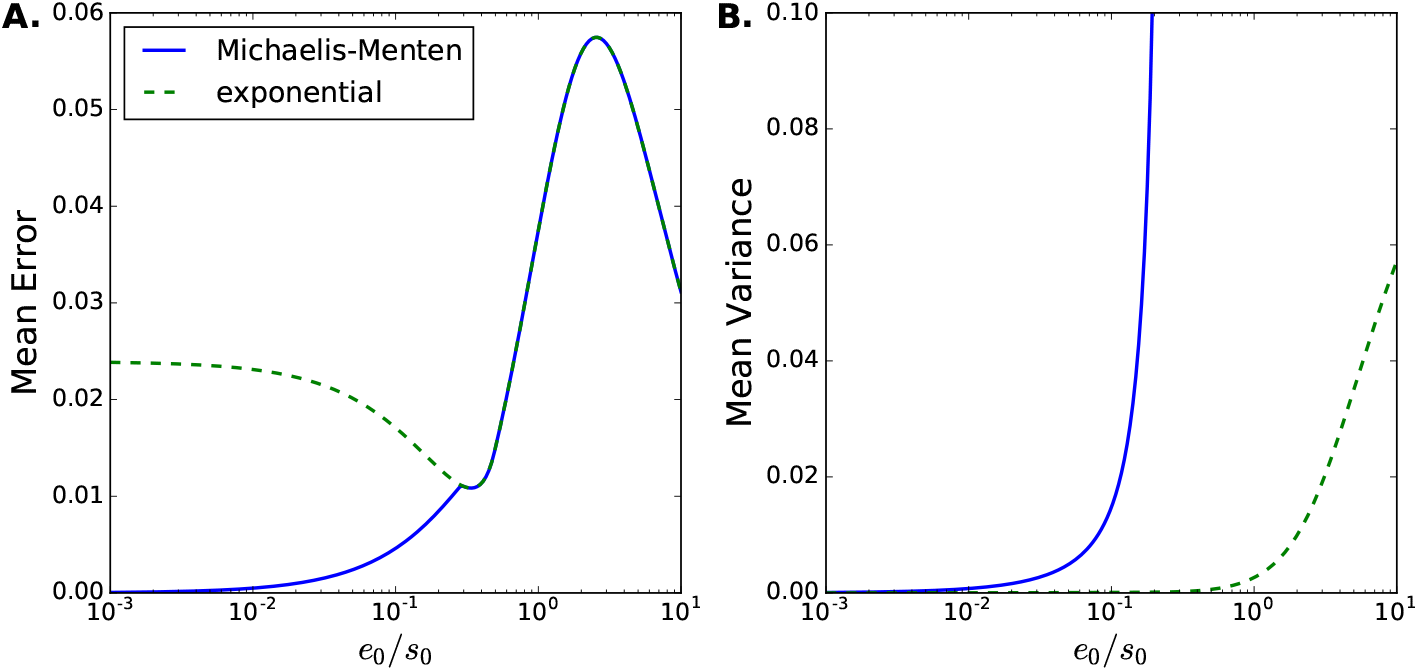
**(A) Mean concentration error,** and **(B) Mean variance in estimated parameters for the Michaelis-Menten equation and an exponential model.** For initial enzyme concentrations smaller than initial substrate concentrations, the Michaelis-Menten equation provides a noticeably better approximation of the true progress curve than does the exponential model, allowing for both *V* and *K_M_* to be uniquely determined. Parameters for the case shown are: (*k*_1_, *k*_−1_, *k*_cat_) = (1.0,0.5, 0.5), *t*_obs_ = 3*t_s_*, *ω* = *t*_obs_/1000, *s*_0_ = 1.

### 4.3. Fitting the initial substrate concentration does not significantly alter estimates of *K_M_ and* V

Even when the reactant stationary assumption is valid, a small amount of substrate will be consumed in the initial transient period. Hence, the substrate concentration at the start of the reaction may not exactly correspond to that at the start of the quasi-stead-state phase. Although this difference is small, it is not clear whether this can noticeably alter the estimation of *K_M_* and *V*. Additionally, time course measurements often employ optical techniques to collect concentration data. Without time consuming calibration curve experiments to relate the fluorescent intensity to concentration directly, only relative concentrations are known. For these reasons, *s*_0_ can be treated as an additional unknown parameter for the regression analysis [42].

**Fig. 5** shows error contours for estimates of *K_M_*, *V* and *s*_0_ for different experimental conditions. Similar to when *s*_0_ is assumed known, the errors in *K_M_* and *V* follow lines of constant *e*_0_/*s*_0_ at low substrate concentration. Additionally, **Fig. 5C** shows that the best-fit value of *s*_0_ corresponds to the true value of the initial substrate concentration for conditions where the reactant stationary assumption is valid. These results indicate that including *s*_0_ as a free parameter can yield similar information about the constants *K_M_* and *V*, even in those cases when no definite concentrations are known.

**Figure 5:**
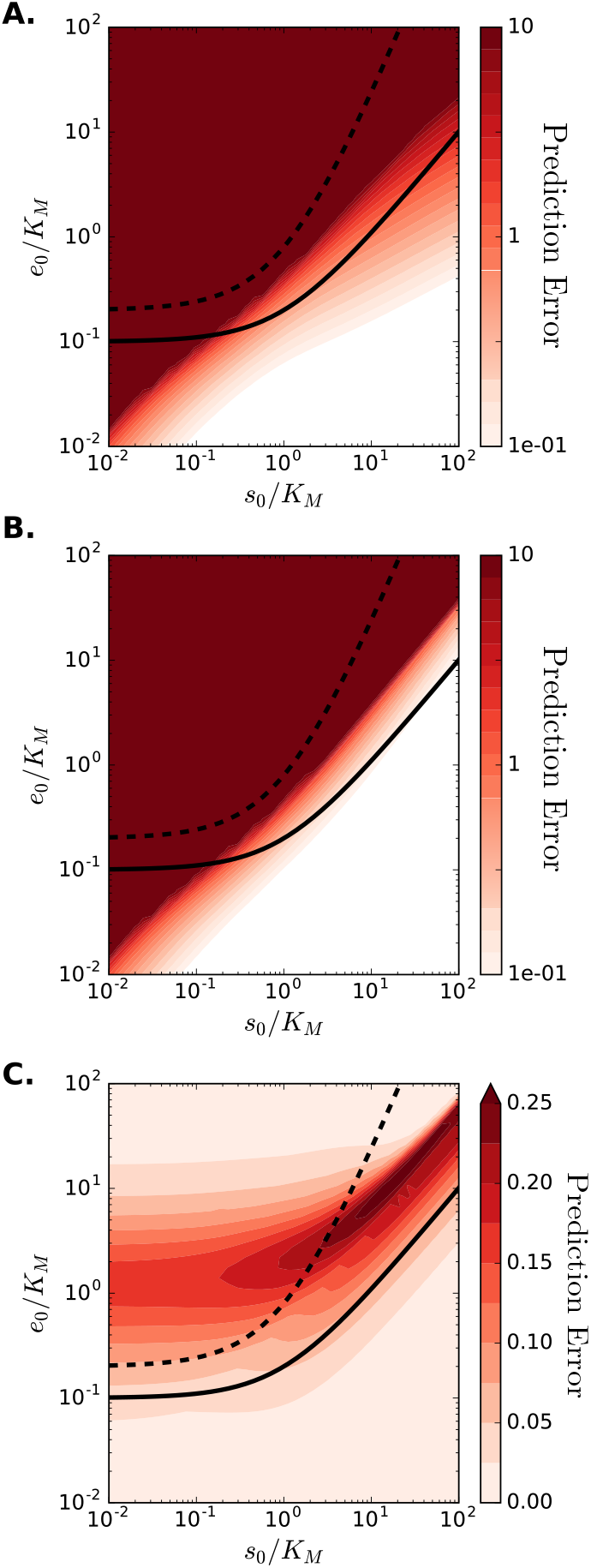
Error contours when initial substrate concentration, *s*_0_, is estimated from data. *K_M_* and *V* prediction errors (panels **A** and **B**, respectively) are qualitatively the same as those found when *s*_0_ is known a priori. The error contours in estimating *s*_0_ (panel **C**) follows the reactant stationary condition, and show accurate estimation is possible when initial enzyme concentration is high and initial substrate concentration is low. Parameters for the case shown are: (*k*_1_, *k*_−1_, *k*_cat_) = (1.0,0.5, 0.5), *t*_obs_ = 3*t_s_*, *ω* = *t*_obs_/100.

### 4.4. Data noise further reduces the range of conditions providing accurate estimates of *K_M_* and *V*

In any physical experiment, some finite amount of measurement error will be present. To understand how signal noise affects the estimation of *K_M_* and *V*, we add noise to the numerically-calculated solution of (5a) such that the data becomes

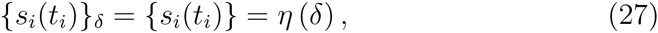

where *η* is a pseudo-random number drawn from a Gaussian distribution of mean zero and standard deviation *δ*. The data is then fitted as described in Section 4.1. However, the noise in the data precludes the use of the method described for estimating good initial conditions for the solver. Without a smoothing procedure, the difference formulas (22) and (23) can lead to large errors. In order to eliminate possible uncertainty arising from the determination of good initial guesses from experimental data, we chose the “true” values of *K_M_* and *V* as the starting point for the optimization algorithm.

Contour plots of the errors in the estimated values of *K_M_* and *V* for the case of *δ* = 0.01 are shown in **Fig. 6.** Qualitatively, they exhibit the same behavior as the noise-free error contours (**Fig. 3**) and display a negligible increase in the magnitude of the error. However, a meaningful characterization of the quality of a fit is the variance in the estimated model parameters. To calculate the variance of *K_M_* and *V*, the covariance matrix is first calculated as

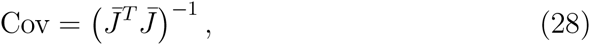

where 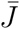 is the Jacobian evaluated numerically at the terminal point of the optimization. The variance for *K_M_* and *V* are then the diagonal elements of Cov. As shown in **Fig. 7**, the range of experimental conditions leading to precise estimates of *K_M_* becomes significantly constrained when even a small amount of measurement error is present. Only in the region where *s*_0_/*K_M_* ≈ 1 and *e*_0_/*K_M_* ≪ 1 are the estimated *K_M_* values robust. At larger initial substrate concentrations, the noise in the data sufficiently smears the sharply curved region of the substrate progress curve, making extraction of *K_M_* prone to uncertainty. At small initial substrate concentrations, the added noise reduces the distinction between the exponential and MM solution branches shown in **Fig. 4A**, making independent determination of *K_M_* and *V* more difficult. Hence, even with only slight measurement error the reliability of estimated parameters falls significantly as the ratio *s*_0_/*K_M_* departs from unity.

**Figure 6:**
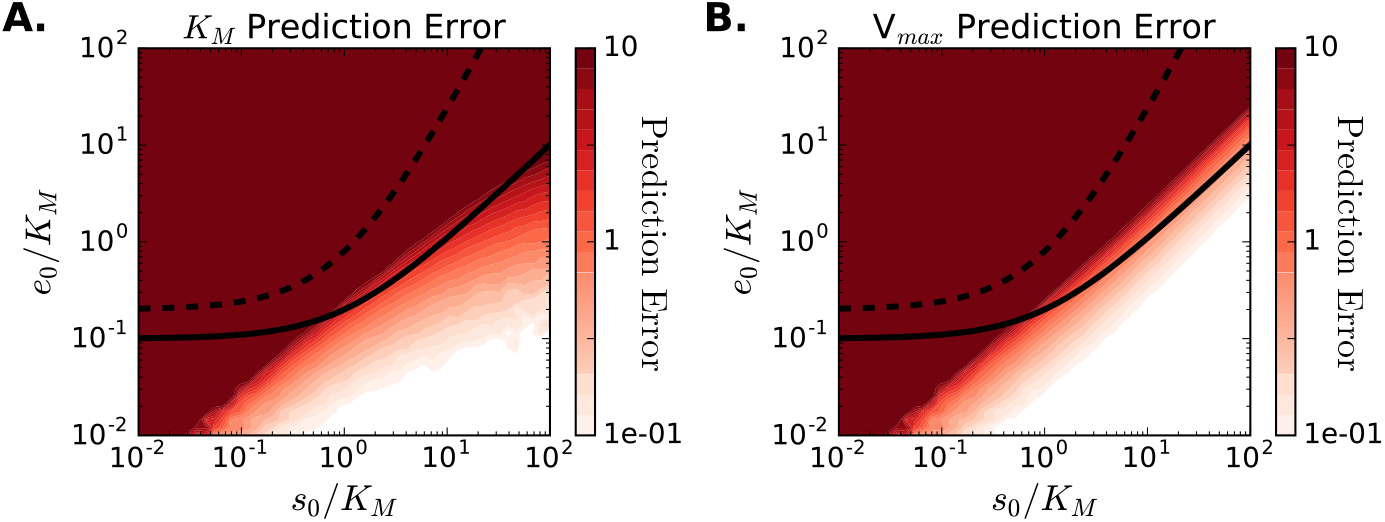
Error contours for data with Gaussian noise. When noise is added to the simulated data (*δ* = 0.01), errors in the estimated parameters worsen compared to noise-free data. Parameters for the case shown are: (*k*_1_, *k*_−1_, *k*_cat_) = (1.0,0.5, 0.5), *t*_obs_ = 3*t_s_*, *ω* = *t*_obs_/1000.

**Figure 7:**
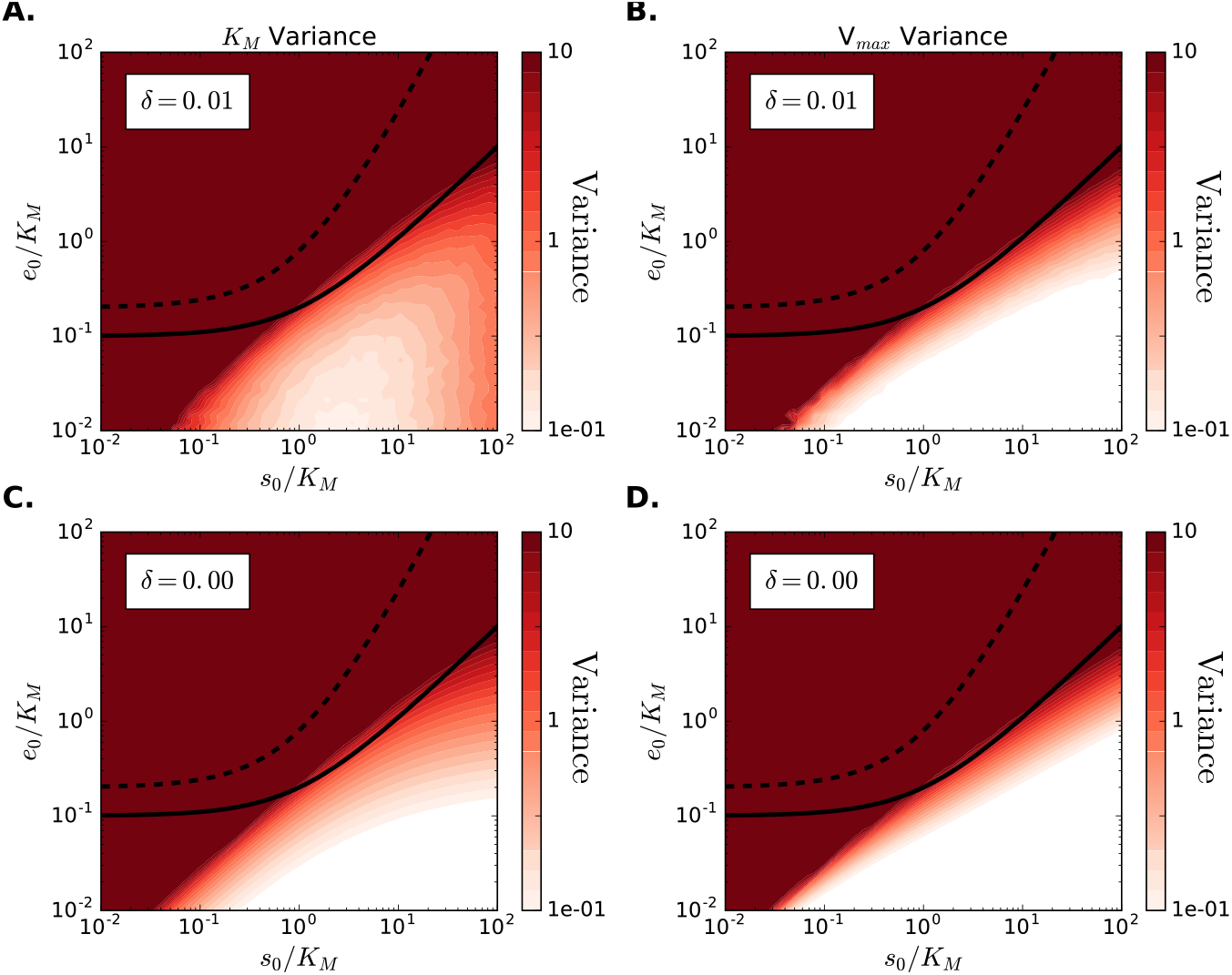
Computed parameter variance for noisy and noise-free data. Estimated variance in the parameters *K_M_* (Panels **A** and **C**) and *V* (Panels **B** and **D**) for cases with *δ* = 0.01 (**A** and **B**) and no noise (**C** and **D**). Even a small amount of noise restricts the range of conditions providing robust parameter estimates. Parameters for the case shown are: (*k*_1_, *k*_−1_, *k*_cat_) = (1.0, 0.5, 0.5), *t*_obs_ = 3*t_s_*, *ω* = tobs/1000.

## 5. Discussion

In this work, we have carried out a systematic analysis of the forward and inverse problems of the MM equation for the single substrate, single enzyme catalyzed reaction. For the forward problem, it is widely believe that the MM equation accurately captures the kinetics of simple enzyme-catalyzed reactions when the reactant-stationary assumption holds true. Through a concentration error analysis, we find that satisfying the reactant-stationary assumption is a sufficient condition for the validity of the MM equation to describe the time course of the enzyme catalyzed reaction. However, the MM equation can accurately describe the reaction dynamics, even when the reactant-stationary assumption is invalid when *K_S_*/*K* ≪ 1 and *s*_0_/*K_M_* ≫ 1 (see, **Fig. 1A**).

As we have shown in this paper, the validity of the MM equation to describe the dynamics of the enzyme catalyzed reaction does not imply that *K_M_* and *V* can be obtained from experimental progress curves conducted within the parameter constraints established by the reactant stationary assumption. This highlights an important problem encountered in parameter estimation. Even when the MM equation very accurately fits the experimental data, the fitted parameters may not accurately represent their true values. Without a thorough analysis of the inverse problem, it is not possible to distinguish between good fits that provide poor parameter estimates, and good fits that accurately estimate parameters.

Most of the research done on the analysis of enzyme progress curves has focused on the nonlinear regression analysis and algorithms to fit progress curve data [43, 35, 28, 44]. Additional research has investigated the design of progress curve experiments from a computational and theoretical standpoint [17]. In these works, either experimental data is collected, or artificial data is generated by adding noise to numerical solutions the integrated MM equation for prescribed values of *K_M_* and *V*. Then, the artificial data is fitted in order to estimate *K_M_* and *V*. Although this procedure can identify values of *K_M_* and *V* for which progress curves can be well-fit by the integrated MM equation, it makes no connection to the underlying microscopic rate constants. Hence, these studies do not directly assess whether the predicted values of *K_M_* and *V* are connected to their microscopic definitions. In the present study we have addressed this issue through two approaches. We first considered the asymptotic behavior of the MM equation under distinct experimental conditions (Section 3). Then, we extracted data from numerical solutions to the underlying mass-action system for prescribed microscopic rate constants, comparing the predicted values of *K_M_* and *V* with those derived from the prescribed values of *k*_1_, *k*_−1_, and *k_cat_*.

The detailed error analysis presented in Section 4 provides guidelines for the ranges of experimental conditions allowing for true parameter estimation. Specifically, we see that, in order for both *K_M_* and *V* to be derived from substrate progress curve measurements:

1. The initial substrate concentration must be within approximately an order of magnitude of the Michaelis constant, that is *s*_0_ = *O*(*K_M_*), especially when significant noise is present in the data. When the initial substrate concentration is in great excess of the Michaelis constant, that is *s*_0_ ≫ *K_M_*, a linear fit to the initial velocity will yield *V*, but provide no information about *K_M_*. When the initial substrate concentration is small compared to the Michaelis constant, that is *s*_0_ ≪ *K_M_*, an exponential fit to the progress curve will provide an estimate for the ratio of *V* to *K_M_*, but neither parameter independently.
2. The initial enzyme concentration must be smaller than the Michaelis constant, that is *e*_0_/*K_M_* ≪ 1, especially when significant noise is present in the data.
3. Data points should be collected around the time point where the time course curvature is at it highest. The length of the high-curvature region is quantified through the timescale *t_Q_* = 27*K_M_s*_0_/4*V*(*K_M_* + *s*_0_). *t_Q_* must not be significantly smaller than *t_S_* if both *K_M_* and *V* are to be estimated from a single progress curve. Theoretically, any two points could be used to estimate *K_M_* and *V*, but empirical statistical analysis carried out elsewhere [14] shows that a minimum of 12 points is ideal for nonlinear regression analysis. These points should sample the region around the point of maximum curvature defined by *t_Q_* (see, **Fig. 2A**).

The above points address important questions necessary to design experiments: What initial substrate concentrations should be used? What initial enzyme concentration should be used? At what time point should data be collected? How many data points should be collected along the curve?

Interestingly, only the first recommendation coincides with previous analysis done by Duggleby and Clarke [17], who recommend an initial substrate concentration of approximately 2.5*K_M_*. However, we additionally provide error contours for parameters estimated from experiments conducted under conditions far from this optimal value. This analysis shows that reasonable estimates can be expected so long as the initial substrate concentration is within an order of magnitude of the optimal value, that is 0.25–25 *K_M_*. Furthermore, noise in the data restricts this range to be significantly smaller than the theoretical range for the validity of the MM equation.

The current experimental practice for data collection is that measurements should be made until the extent of the reaction reaches 90%. Duggleby and Clarke [17] finds that there is no advantage of extending beyond this point. In our analysis, we discovered that errors in *K_M_* and *V* are minimized when data is collected in the region around the point of maximum curvature defined by *t_Q_* (see, Figure 2A).

In general, since these requirements listed above depend on *K_M_*, they cannot be assessed before conducting an experiment. However, they do provide useful checks that can reduce the number of experiments required, especially when compared to parameter estimation based on initial rate experiments. If a progress curve for a given initial substrate concentration cannot be fitted by an exponential, and has a curvature that can be resolved, nonlinear regression of the progress curve will provide accurate estimates of both *K_M_* and *V*. If, say, the progress curve can reasonably be fit by an exponential, a second experiment with substantially larger initial substrate concentration should be performed. Then, the second progress curve, so long as the increase in initial substrate concentration is great enough to surpass the substrate range for with the kinetics are exponential, should yield either a curve from which both parameters can be estimated, or a curve from which V can be estimated. Hence, the two experiments are sufficient to make preliminary estimates of both *K_M_* and *V*. This is in contrast to initial rate experiments, which require a large number of experiments such that a curve of the initial reaction velocity as a function of s0 can be produced. For accurate measurement of *K_M_* and *V* from initial rate experiments, both large and small values of the substrate (relative to *K_M_*) must be used [14, 31]. Hence, progress curve analysis will always require fewer experiments than initial rate experiments. Additionally, if initial rate experiments are used, progress curve analysis can be used as a check the accuracy of the estimates. Values of *K_M_* and *V* obtained from fitting (5a) to the initial rate data should correspond to those values obtained from progress curve analysis of the experiments for which the initial rates are intermediate between 0 and *V*.

In conclusion, this work both advocates and cautions the use of progress curve analysis in modeling and determining kinetic parameters for enzymatic reactions. Progress curve assays can greatly reduce the number of experiments (and hence the cost and quantity of reagents) needed, while still providing accurate measurements. However, it is essential to not conflate an accurate fit with an accurate estimate of *K_M_* and *V*. If this is kept in mind, progress curve analysis has significant advantages over the use of initial rate experiments.

## Acknowledgments

This work is supported by the University of Michigan Protein Folding Diseases Initiative.

